## Supplementary Material for "Aberration correction in long GRIN lens-based microendoscopes for extended field-of-view two-photon imaging in deep brain regions"

**Figure supplements**

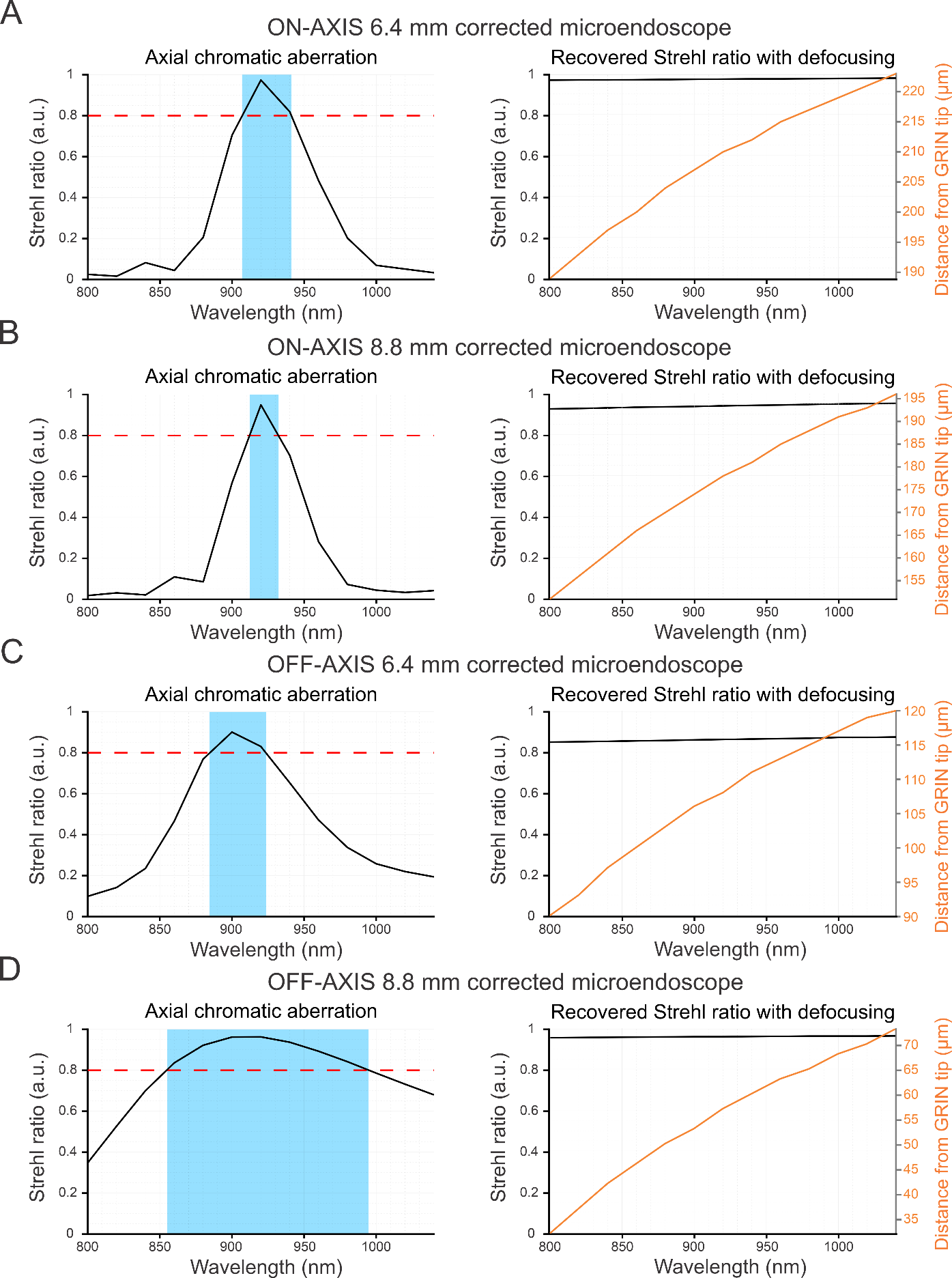

**Figure 1-figure supplement 1. Performance of corrected microendoscopes at different wavelengths.** A) Left: simulated Strehl ratio on the optical axis as a function of wavelength for the 6.4 mm-long corrected microendoscope. The red dashed line marks the diffraction-limited threshold according to the Maréchal criterion. The area highlighted in light blue indicates the range of wavelengths for which the Strehl ratio is above the diffraction-limited threshold. Right: maximal simulated Strehl ratio obtained on-axis (black line) and corresponding working distance (Distance from GRIN tip, orange line) as a function of wavelength. B) Same as in (A) for the 8.8. mm-long corrected microendoscope. C) Same as in A) for Strehl ratio computed off-axis, considering the largest simulated radial distance from the optical axis used to determine the profile of the corrective lens. In the object plane, this off-axis radial distance is equal to 156 μm. D) Same as in C) for the 8.8. mm-long corrected microendoscope and largest simulated radial distance equal to 135 μm.

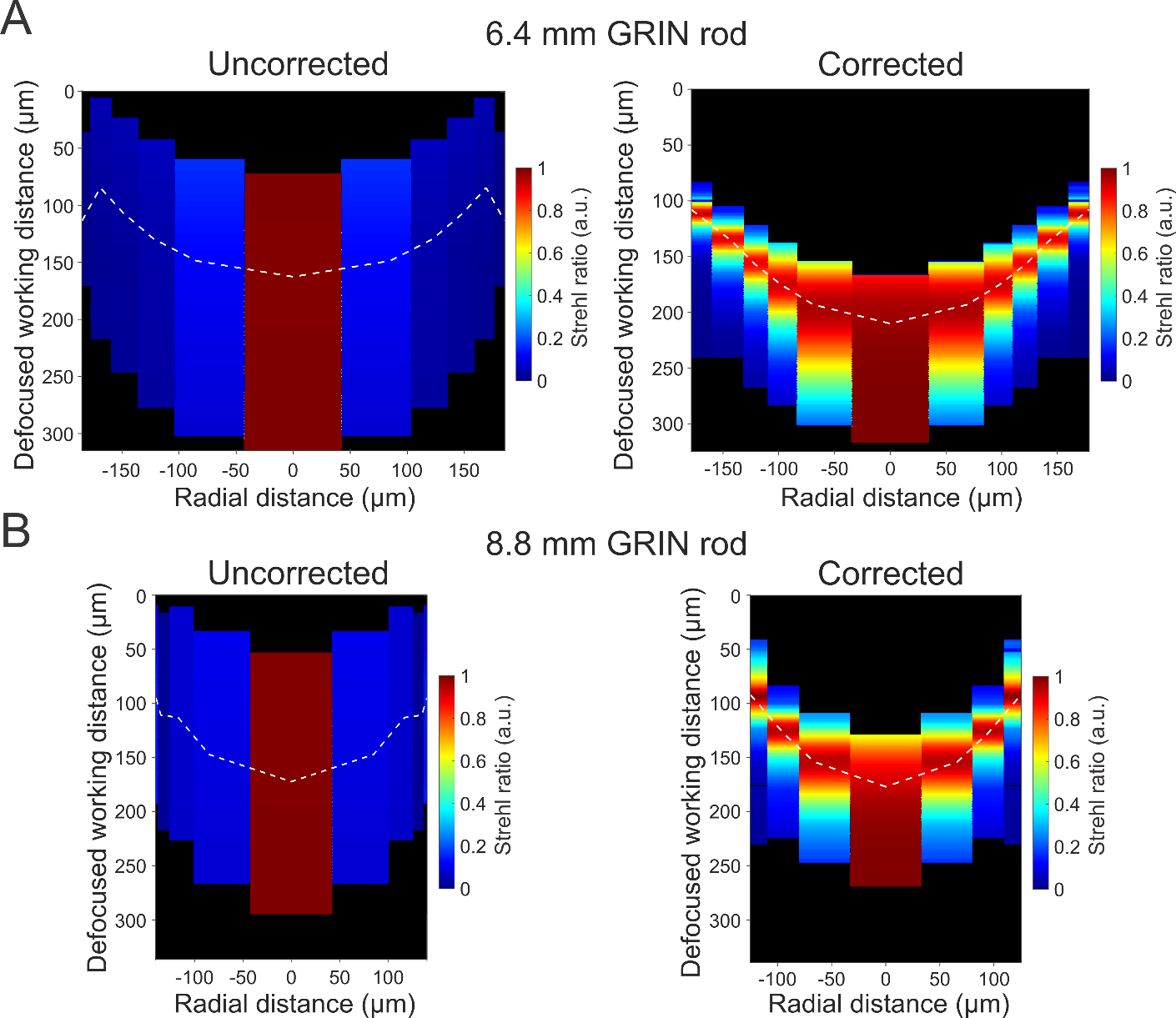

**Figure 1-figure supplement 2. Optical performance in out-of-focus planes.** A) Pseudocolor map of the Strehl ratio obtained in the object space (*x,*z projection) at different focal distances (*z*-axis, Defocused working distance) and as a function of radial distance from the optical axis (corresponding to the zero in the *x*-axis) computed on the focal plane in the object space for the uncorrected (left) and the corrected (right) 6.4 mm-long microendoscope. For the corrected microendoscope (right), the white dashed line represents the focal plane for which the corrective lens profile was optimized. For the uncorrected microendoscope (left), the white dashed line is the focal plane of the system in the absence of the corrective lens. B) Same as in (A) for the 8.8 mm-long microendoscope.

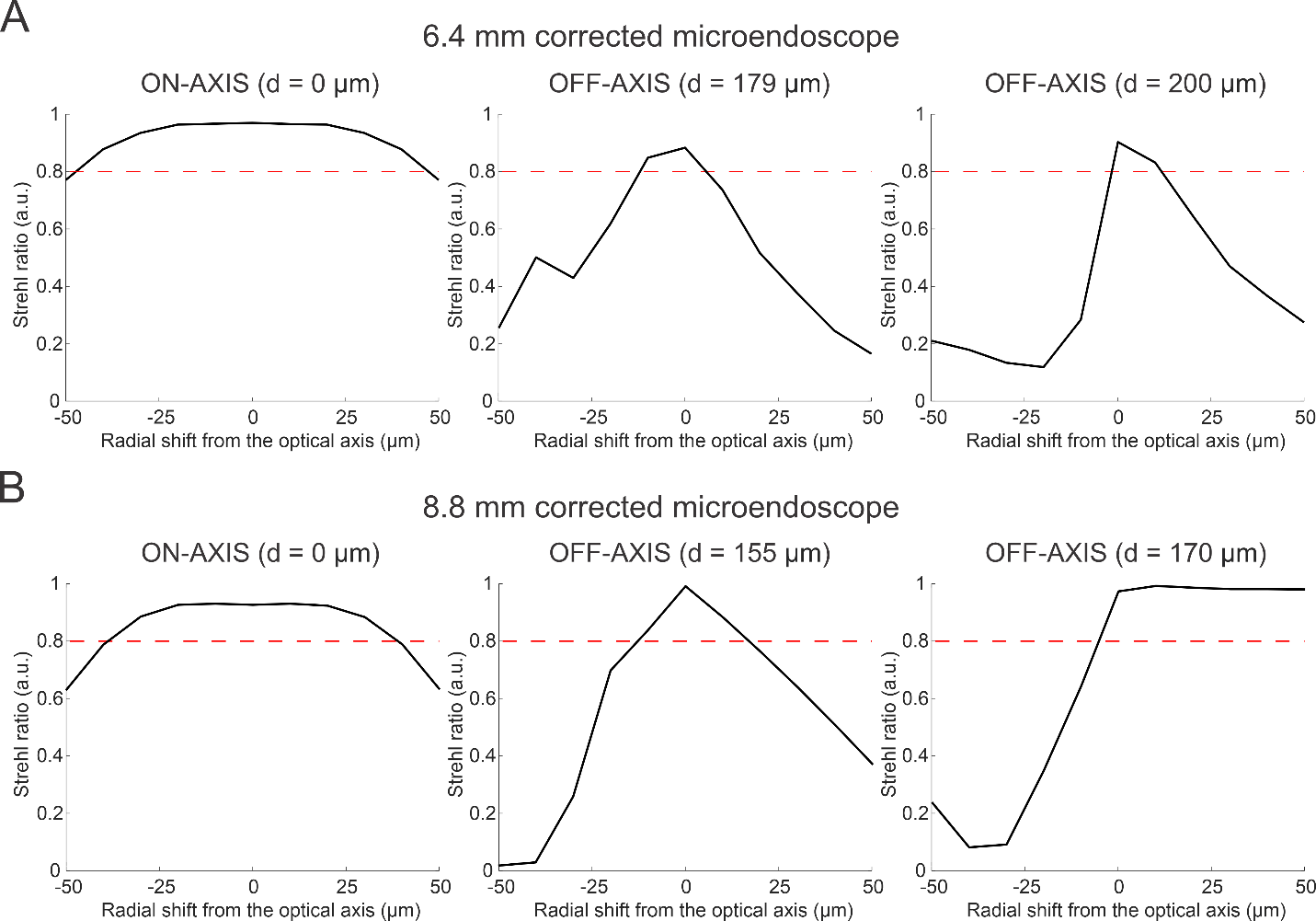

**Figure 1-figure supplement 3. Optical performance of corrected microendoscopes as a function of decentering the corrective lens.** A) Strehl ratio as a function of radial shift between the corrective lens and the optical axis of the GRIN rod for the 6.4 mm-long corrected microendoscope. Simulation were performed along the optical axis of the GRIN rod (left, on-axis, d = 0 µm) and at two marginal radial distances (d, center and right panels; radial distances are computed on the focal plane in the image space, before the GRIN rod). The red dashed line marks the diffraction-limited threshold according to the Maréchal criterion. B) Same as in (A) for the 8.8 mm-long corrected microendoscope.

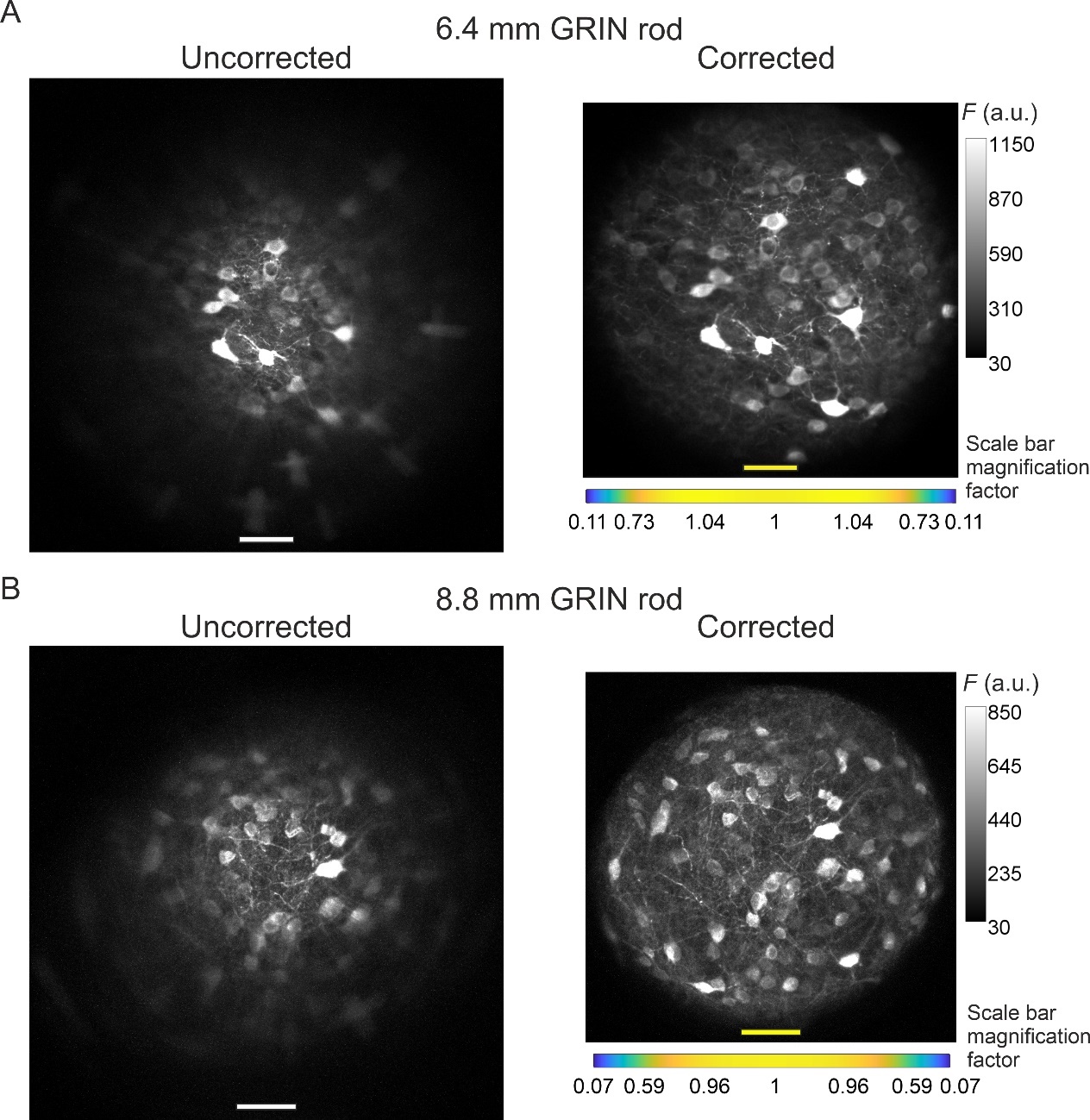

**Figure 4-figure supplement 1. Modified version of Figure 4 with the FOVs of corrected microendoscopes rescaled to match the real pixel size of the FOVs of uncorrected microendoscopes in the center of the image.** See caption of Figure 4. In Figure 4A, B, images acquired with corrected microendoscopes (right images) have a real pixel size in the center of the FOV smaller than the pixel size in the center of the FOVs of uncorrected microendoscopes (left images), due to distortion introduced by corrective lenses (Figure 3E, F). Here, the FOV of corrected microendoscopes (right images) are scaled in order to have the scale bar in the center of the FOV equal to the one of the FOV of uncorrected microendoscopes (left images).

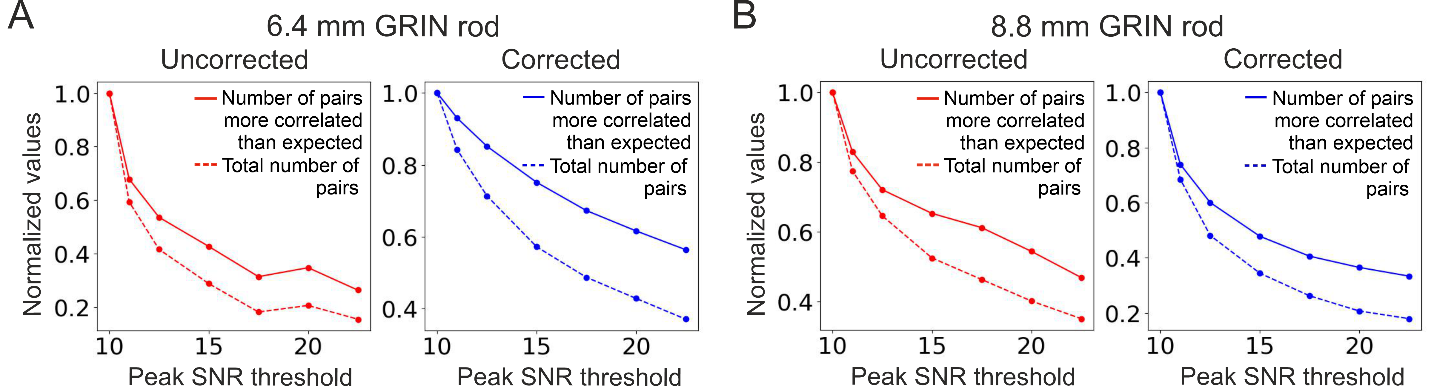

**Figure 6-figure supplement 1. The total number of detected adjacent cell pairs decreases faster with peak SNR threshold than the number of adjacent cell pairs more correlated than expected does.** A) Number of adjacent cell pairs more correlated than expected (solid lines) and total number of detected adjacent cell pairs (dashed lines) as a function of the peak SNR threshold for the uncorrected (red, left) and the corrected (blue, right) 6.4 mm-long microendoscope (see also Figure 6A). Values are normalized to their value at peak SNR threshold = 10. B) Same as (A) for the 8.8 mm-long microendoscope (see also Figure 6F).

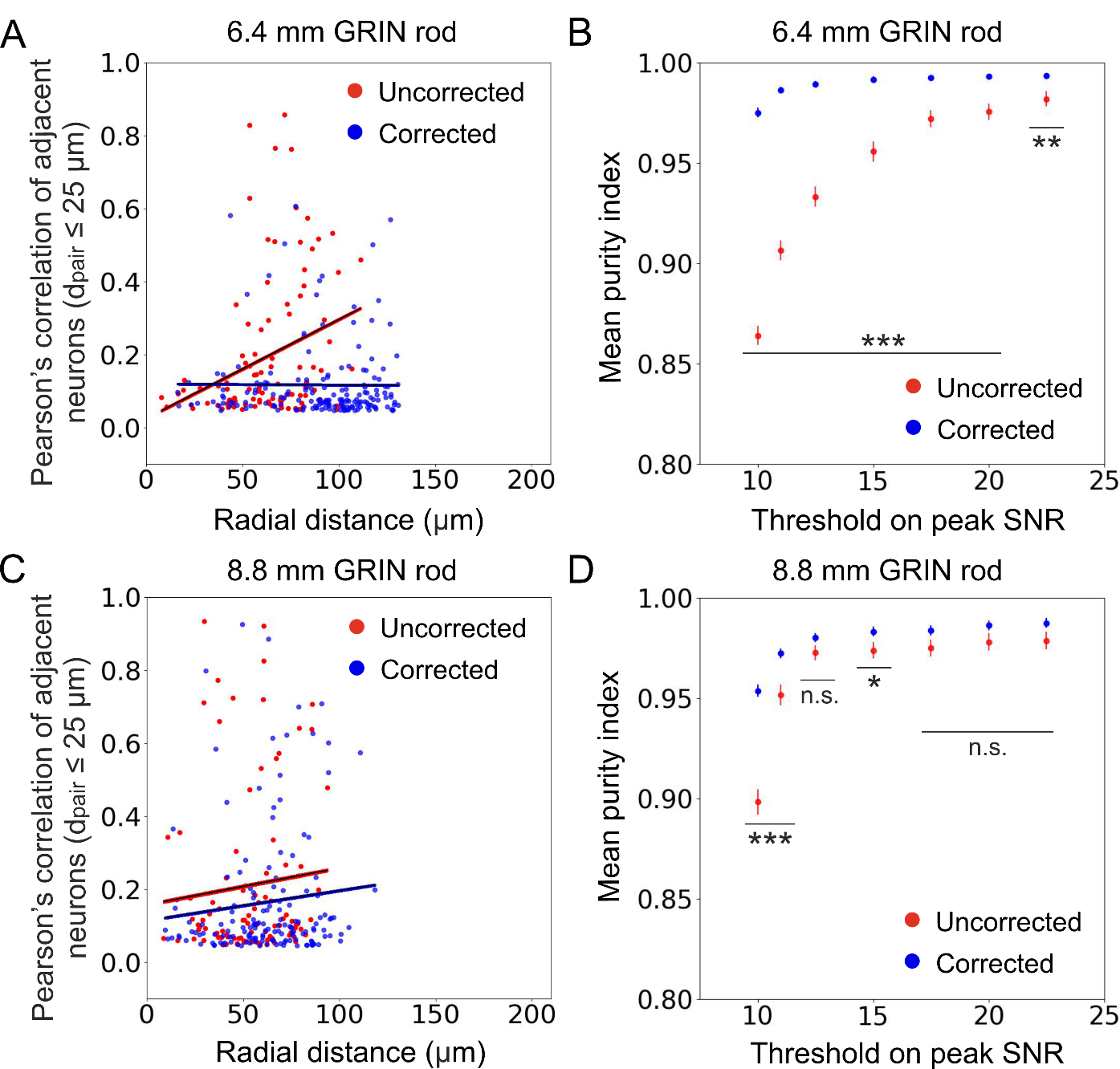

**Figure 6-figure supplement 2. Aberration correction enables more accurate measurement of population activity.** A) Pearson’s correlation of adjacent cell pairs as a function of the radial distance from the center of the FOV in simulated calcium data. Pairs of cells were defined as adjacent if the distance between detected cell centroids d_pair_ was ≤ 25 µm. Pearson’s correlation is displayed only for cell pairs which are more correlated than expected (expected pair correlation was estimated as mean Pearson’s correlation between ground truth activity traces of any possible neuronal pairs plus 3 SDs, see Supplementary Table 5). Data are displayed only for cells with peak SNR > 15 from *n* = 13 simulated experiments with the 6.4 mm-long uncorrected (red) and corrected (blue) microendoscope. Black lines represent the linear regression of data ± 95% confidence intervals (shaded colored areas). Slopes ± s.e.: uncorrected, (0.003 ± 0.002) μm^-1^, significantly different from zero, *p* = 0.00080, permutation test; corrected, (-0.00003 ± 0.00060) μm^-1^, not significantly different from zero, *p* = 0.88, permutation test. Uncorrected, *n* = 102; corrected, *n* = 172. Statistical comparison of slopes, *p* < 10^-10^, permutation test. B) Mean purity index (see Materials and Methods for definition) ± s.e.m. of extracted traces as a function of the peak SNR threshold for *n* = 13 simulated experiments with the 6.4 mm-long uncorrected (red) and corrected (blue) microendoscope. Statistical differences of the means are assessed with the permutation test; **, *p* < 0.01; ***, *p* < 0.001. C) Same as (A) for *n* = 15 simulated experiments with the 8.8 mm-long uncorrected and corrected microendoscope. Slopes ± s.e.: uncorrected, (0.001 ± 0.002) μm^-1^, not significantly different from zero, *p* = 0.41, permutation test; corrected, (0.0008 ± 0.0013) μm^-1^, not significantly different from zero, *p* = 0.20, permutation test. Uncorrected, *n* = 96; corrected, *n* = 152. Statistical comparison of slopes, *p* = 0.86, permutation test. D) Same as (B) for *n* = 15 simulated experiments with the 8.8 mm-long uncorrected and corrected microendoscope. *, *p* < 0.05; ***, *p* < 0.001; n.s., not significant.

**Supplementary files**

|  | ***c*** | ***k*** | ***α_1_*** | ***α_2_*** | ***α_3_*** | ***α_4_*** |
| --- | --- | --- | --- | --- | --- | --- |
| **6.4 mm-long GRIN rod** | 0.25 | 0 | -4.00 | 2.65 | 3.53 | -1.57 |
| **8.8 mm-long GRIN rod** | -0.36 | 0 | -0.94 | 23.40 | 349.33 | -720.06 |

**Supplementary Table 1. Parameters of the polynomial function describing the aspherical surface of simulated corrective lenses.** Parameters of equation (*1*) (see Materials and Methods) for the simulated corrective lens designed to be applied to the GRIN rods of length 6.4 mm (top row) and 8.8 mm (bottom row).

| **Microendoscope based on 6.4 mm-long GRIN rod** | | | | | |
| --- | --- | --- | --- | --- | --- |
| **Uncorrected** | | | **Corrected** | | |
| **Radial distance (µm)** | **Axial resolution (µm)** | **Lateral resolution (µm)** | **Radial distance (µm)** | **Axial resolution (µm)** | **Lateral resolution (µm)** |
| 0 | 12.7 | 1.6 | 0 | 7.3 | 0.7 |
| 89 | 39.4 | 2.7 | 65 | 7.3 | 0.7 |
| 127 | 29.5 | 14.4 | 93 | 9.0 | 1.2 |
| 155 | 29.9 | 19.0 | 115 | 11.2 | 1.2 |
| 179 | 31.6 | 23.4 | 135 | 13.7 | 1.3 |
| 200 | 24.5 | 24.9 | 155 | 19.9 | 1.6 |
| **Microendoscope based on 8.8 mm-long GRIN rod** | | | | | |
| **Uncorrected** | | | **Corrected** | | |
| **Radial distance (µm)** | **Axial resolution (µm)** | **Lateral resolution (µm)** | **Radial distance (µm)** | **Axial resolution (µm)** | **Lateral resolution (µm)** |
| 0 | 13.5 | 1.7 | 0 | 7.5 | 1.2 |
| 88 | 24.2 | 3.7 | 63 | 9.0 | 0.8 |
| 125 | 30.8 | 19.9 | 91 | 13.6 | 1.2 |
| 152 | 29.8 | 22.7 | 116 | 24.2 | 1.9 |

**Supplementary Table 2. Spatial resolution of simulated uncorrected and corrected microendoscopes.** Axial and lateral resolution of simulated microendoscopes were evaluated measuring the dimensions of simulated 2P PSF for each probe at different radial distances. *x,z* (Axial) and *x,y* (Lateral) intensity profiles of simulated PSFs were fitted with Gaussian curves and their FWHM was used to define the resolution, as done for experimental PSFs (see Materials and Methods).

| **Local magnification factor** | | | | | |
| --- | --- | --- | --- | --- | --- |
| **Microendoscope based on 6.4 mm-long GRIN rod**  *f(x) = ax^4^ + bx^2^ + c* | | | **Microendoscope based on 8.8 mm-long GRIN rod**  *f'(x) = a'x^4^ + b'x^2^ + c'* | | |
|  | **Uncorrected** | **Corrected** |  | **Uncorrected** | **Corrected** |
| *a* | -0.23∙10^-9^  (-1.12∙10^-9^, 0.66∙10^-9^) | -0.83∙10^-9^  (-1.09∙10^-9^,  -0.58∙10^-9^) | *a'* | 0.52 ∙10^-9^  (-1.52∙10^-9^, 2.56∙10^-9^) | -0.28∙10^-9^  (-0.56∙10^-9^, 0.0059∙10^-9^) |
| *b* | 0.15∙10^-5^  (-1.07∙10^-5^, 1.37∙10^-5^) | 0.10∙10^-4^  (0.036 ∙10^-4^, 0.17∙10^-4^) | *b'* | -0.27∙10^-5^  (-2.63∙10^-5^, 2.08∙10^-5^) | -0.18∙10^-4^  (-0.26 ∙10^-4^,  -1.07∙10^-4^) |
| *c* | 1.03  (1.00, 1.06) | 1.29  (1.25, 1.32) | *c'* | 1.04  (0.99, 1.09) | 1.36  (1.32, 1.40) |
| R-square | 0.12 | 0.93 | R-square | 0.13 | 0.98 |
| **Radial distance calibration** | | | | | |
| **Microendoscope based on 6.4 mm-long GRIN rod**  *g(x) = dx^4^ + ex^2^ + fx + h* | | | **Microendoscope based on 8.8 mm-long GRIN rod**  *g'(x) = d'x^4^ + e'x^2^ + f'x +h'* | | |
|  | **Uncorrected** | **Corrected** |  | **Uncorrected** | **Corrected** |
| *d* | -0.14∙10^-7^  (-0.31∙10^-7^, 0.33∙10^-7^) | -0.33∙10^-7^  (-0.40∙10^-7^,  -0.27∙10^-7^) | *d'* | 0.30∙10^-8^  (-2.28∙10^-8^, 2.88∙10^-8^) | -0.42∙10^-7^  (-0.46∙10^-7^,  -0.38∙10^-7^) |
| *e* | 0.22∙10^-3^  (-0.24∙10^-3^, 0.68∙10^-3^) | 1.03∙10^-3^  (0.68∙10^-3^, 1.37∙10^-3^) | *e'* | 0.45∙10^-4^  (-5.25∙10^-4^, 6.16∙10^-4^) | -0.16∙10^-4^  (-2.44∙10^-4^, 2.11∙10^-4^) |
| *f* | 1.03  (1.00, 1.06) | 1.24  (1.20, 1.27) | *f'* | 1.04  (1.00, 1.08) | 1.35  (1.32, 1.37) |
| *h* | -0.17  (-0.67, 0.33) | 0.13  (-0.63, 0.89) | *h'* | -0.18  (-0.70, 0.34) | -0.37  (-0.87, 0.14) |
| R-square | 1.00 | 1.00 | R-square | 1.00 | 1.00 |

**Supplementary Table 3. Parameters used for the computation of the local pixel size and for distance calibration of images acquired with microendoscopes.** Coefficients of the quartic functions fitting the measurements performed on images acquired on the calibration ruler for uncorrected and corrected microendoscopes based on the 6.4 mm-long GRIN rod (left) and the 8.8 mm-long GRIN rod (right). The numbers in parenthesis indicate the 95% lower and upper confidence bounds (see Figure 3E, F). R-square values are indicated for each fit.

| **Axial resolution** | | | | | |
| --- | --- | --- | --- | --- | --- |
| **Microendoscope based on 6.4 mm-long GRIN rod**  *f(x) = ax^4^ + bx^2^ + c* | | | **Microendoscope based on 8.8 mm-long GRIN rod**  *f'(x) = a'x^4^ + b'x^2^ + c'* | | |
|  | **Uncorrected** | **Corrected** |  | **Uncorrected** | **Corrected** |
| *a* | 0.41∙10^-6^  (-0.39∙10^-6^, 1.21∙10^-6^) | 0.20∙10^-7^  (-0.24∙10^-7^,  0.65 ∙10^-7^) | *a'* | 0.31∙10^-6^  (-1.28∙10^-6^, 1.91∙10^-6^) | -0.31∙10^-7^  (-0.63 ∙10^-7^, 0.014∙10^-7^) |
| *b* | 0.59∙10^-4^  (-43.64∙10^-4^, 44.83∙10^-4^) | -0.52∙10^-4^  (-8.53∙10^-4^, 7.48∙10^-4^) | *b'* | 0.19∙10^-2^  (-0.68∙10^-2^, 1.05∙10^-2^) | 0.12∙10^-2^  (0.053∙10^-2^, 0.18∙10^-2^) |
| *c* | 7.99  (4.20, 11.77) | 8.52  (6.03, 11.00) | *c'* | 7.43  (0.089, 14.78) | 7.04  (4.77, 9.30) |
| R-square | 1.00 | 0.72 | R-square | 1.00 | 0.93 |
| **Lateral resolution** | | | | | |
| **Microendoscope based on 6.4 mm-long GRIN rod**  *g(x) = dx^4^ + ex^2^ + f* | | | **Microendoscope based on 8.8 mm-long GRIN rod**  *g'(x) = d'x^4^ + e'x^2^ + f'* | | |
|  | **Uncorrected** | **Corrected** |  | **Uncorrected** | **Corrected** |
| *d* | 0.26∙10^-7^  (-2.83∙10^-7^, 3.35∙10^-7^) | 0.11∙10^-7^  (0.050∙10^-7^, 0.16∙10^-7^) | *d'* | -0.71∙10^-8^  (-22.84∙10^-8^, 21.42∙10^-8^) | 0.22∙10^-8^  (-0.047∙10^-8^, 0.50∙10^-8^) |
| *e* | 0.26∙10^-3^  (-1.45∙10^-3^, 1.97∙10^-3^) | -0.85∙10^-4^  (-1.87∙10^-4^, 0.17∙10^-4^) | *e'* | 0.13∙10^-3^  (-1.10∙10^-3^, 1.35∙10^-3^) | 0.59∙10^-4^  (0.031∙10^-4^, 1.15∙10^-4^) |
| *f* | 1.10  (-0.36, 2.56) | 1.37  (1.06, 1.69) | *f'* | 1.06  (0.0089, 2.10) | 1.12  (0.93, 1.31) |
| R-square | 0.99 | 0.95 | R-square | 0.94 | 0.98 |

**Supplementary Table 4. Fitting parameters for PSF measurements of uncorrected and corrected microendoscopes.** Coefficients of quartic functions fitting experimental PSF data (axial, top; lateral, bottom) are presented for uncorrected and corrected microendoscopes based on the 6.4 mm-long GRIN rod (left) and the 8.8 mm-long GRIN rod length (right). Parentheses indicate the 95% lower and upper confidence bounds (see Figure 3I, J). R-square values are indicated for each fit.

|  | **6.4 mm-long microendoscope**  (*n* = 13 FOVs) | | | **8.8 mm-long microendoscope**  (*n* = 15 FOVs) | | |
| --- | --- | --- | --- | --- | --- | --- |
|  | **Mean** | **Standard deviation** | **Expected pair correlation** | **Mean** | **Standard deviation** | **Expected pair correlation** |
| **Uncorrected** | 0.0393 | 0.0032 | 0.0488 | 0.0386 | 0.0031 | 0.0479 |
| **Corrected** | 0.0388 | 0.0026 | 0.0466 | 0.0389 | 0.0023 | 0.0458 |

**Supplementary Table 5. Expected Pearson’s correlation of cell pair in synthetic calcium data.** Numerical values used to estimate the expected correlation between cell pairs in synthetic calcium t-series are indicated for each microendoscope type. The table displays the mean and SD of Pearson’s correlation between the activity traces of any possible ground truth source neuron pair obtained from *n* simulated FOVs and the expected cell pair correlation (mean Pearson’s correlation plus three SDs). These parameters were used for the analysis in Figure 6A, F and in Figure 6-figure supplement 2A, C.

|  | **Adjacent neurons: distance between centroids ≤ 25 µm** | | |
| --- | --- | --- | --- |
|  | **jGCaMP8f (4 mice)** | **jGCaMP7f (2 mice)** | **Merge (6 mice)** |
| ***n*** | 168 | 27 | 195 |
| ***p* value (normality of residuals)** | D’Agostino-Pearson: 0.51 | Shapiro-Wilk: 0.74 | D’Agostino-Pearson: 0.29 |
| **slope (µm^-1^)** | -0.0006 ± 0.0004 | 0.002 ± 0.001 | -0.0006 ± 0.0004 |
| ***p* value (slope = 0)** | Wald test: 0.11 | Wald test: 0.18 | Wald test: 0.09 |
|  | **Adjacent neurons: distance between centroids ≤ 30 µm** | | |
|  | **jGCaMP8f (4 mice)** | **jGCaMP7f (2 mice)** | **Merge (6 mice)** |
| ***n*** | 249 | 39 | 288 |
| ***p* value (normality of residuals)** | D’Agostino-Pearson: 0.25 | Shapiro-Wilk: 0.62 | D’Agostino-Pearson: 0.42 |
| **slope (µm^-1^)** | -0.0005 ± 0.0003 | 0.001 ± 0.001 | -0.0005 ± 0.0003 |
| ***p* value (slope = 0)** | Wald test: 0.13 | Wald test: 0.18 | Wald test: 0.06 |

**Supplementary Table 6. Linear regression analysis for pairwise correlation of adjacent neurons as a function of the radial distance of pair centroid for *in vivo* 2P imaging data.** The values of the slope of the linear fits are indicated ± s.e. for adjacent neurons with maximum centroid distance equal to 25 µm (top) or 30 µm (bottom). Results obtained with jGCaMP8f, jGCaMP7f, and with the merged dataset including both jGCaMP8f and jGCaMP7f are displayed. The number *n* of adjacent neuron pairs, the *p* value of the indicated statistical test for the normality of residuals, and the *p* value of the Wald test on the null hypothesis of slope = 0 are indicated for each condition.
